## Supplementary material for "HQAlign: Aligning nanopore reads for SV detection using current-level modeling"

### 1 Methods

#### 1.1 Quantization method from QAlign [1]

The nucleotide sequences are inferred from the nanopore current signals by basecallers, therefore, using a $Q$-mer map to translate the basecalled sequences to the current levels implicitly maintains all of the "equivalent" basecalled sequences that could be inferred from the observed current levels. These current levels can be quantized to an alphabet of finite size.

Mathematically, the quantization process is as follows. Let $\Sigma = \{A, C, G, T\}$ be the alphabet of nucleotide sequences. For a symbol $s \in \Sigma$, let $\bar{s}$ be the Watson-Crick complement of $s$. A string $x = s_1 s_2 \ldots s_n$ over $\Sigma$ is called a nucleotide sequence, where $|x| = n$ is the string length and the reverse complement of $x$ is $\overline{x} = \overline{s_1 s_2 \ldots s_n} = \bar{s}_n \bar{s}_{n-1} \ldots \bar{s}_1$. Let $p(x)$ be a list of all $Q$-mers (e.g. $Q$=6) in the string $x$, sorted by their occurrences. For example, $p(x) = k_1 k_2 \ldots k_{n-Q+1}$ and each $Q$-mer $k_i = s_i s_{i+1} \ldots s_{i+Q-1}$ for $i = 1, 2, \ldots, n - Q + 1$. Now, we define $f : \Sigma^Q \to \mathbb{R}$ as the $Q$-mer map [1], which is a deterministic function that translates each $Q$-mer ($k_i$) to the (median) current level (Figure 1b). Now, let $C(x) = c_1 c_2 \ldots c_{n-Q+1}$ be the sequence of the current levels, so that $c_i = f(k_i)$ for $i = 1, 2, \ldots, n - Q + 1$. The current sequence $C(x)$ can be further quantized into $w(x) = q_1 q_2 \ldots q_{n-Q+1}$ by applying hard thresholding function $q_i = g(c_i)$. The thresholding can be ternary ($q_i \in \{0, 1, 2\}$) for $HQ3$ (Figure 1c and Supplemental Figure 1). We define $w(\overline{x})$ as the quantized reverse complementary of sequence $x$, so $\overline{w}(x) = w(\overline{x})$. Supplemental Figure 1 explains this process using a toy example.

#### 1.2 Generalization of HQAlign method

##### 1.2.1 Initial alignment

The nucleotide query $x$ is aligned to a set of nucleotide target sequences $t = (t_1, t_2, \ldots, t_m)$ using Minimap2. This is similar to aligning a read to a genome which has several chromosome

[1] $Q$-mer map is determined by the chemistry of the nanopore flow cell, and is therefore dataset dependent, *i.e.*, the $Q$-mer map for sequencing using R9 flow cell is different from $Q$-mer map for sequencing using R9.4.1 flow cell. The $Q$-mer maps used in this work are generated by Nanopolish (https://github.com/jts/nanopolish).

sequences. This step identifies the region of interests on the target $t$, say, $t_j[s_i : e_i]$, where $t_j$, $j \in \{1, 2, \ldots, m\}$ represent alignment to one or more target chromosomes that $x$ aligns to, $i \in \{1, 2, 3, \ldots\}$ represent represent one or more alignments to chromosome $j$, $s_i$ and $e_i$ are the corresponding start and end location of each alignment $i$ on the target $t_j$, respectively.

##### 1.2.2 Hybrid alignment

In this step, the query $x$ is re-aligned to an extended region of interest on the target $t_j[s_i^q : e_i^q]$ using the QAlign method, where $s_i^q = s_i - b_i$ and $e_i^q = e_i + b_i$, $b_i = (1 - f_i + 0.25)n$ is an appended extension of the region of interest on target, $f_i = (e_i - s_i)/n$ is the fraction of read aligned in initial step, and $n$ is the length of the query $x$. The nucleotide query $x$ and the nucleotide extended target $t_j[s_i^q : e_i^q]$ are converted to the quantized query $x^q$, quantized reverse complement query $\overline{x}^q$ and quantized extended target $t_j^q[s_i^q : e_i^q]$, respectively, using the quantization method demonstrated in QAlign. These quantized sequences are then aligned using modified minimap2 pipeline.

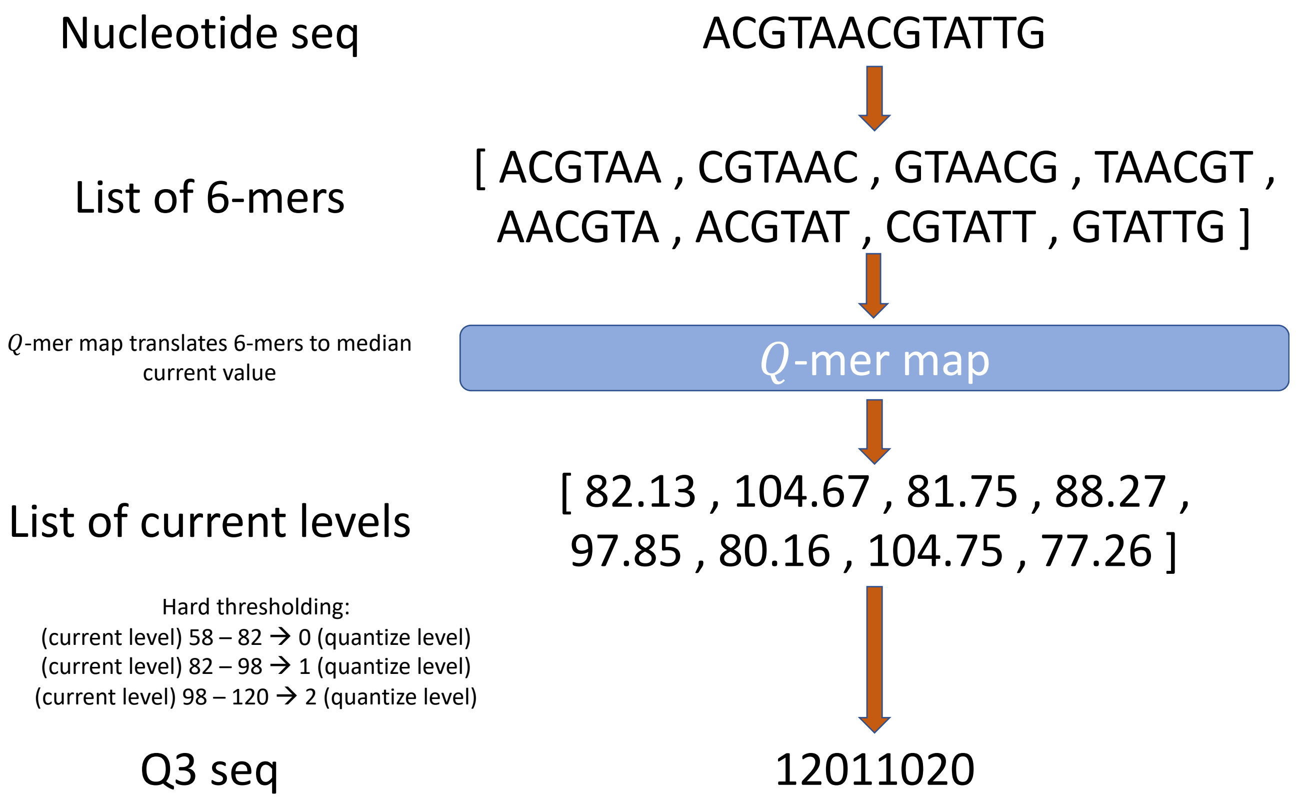


Figure 1: **An example for the Quantization method for QAlign.** The nucleotide sequences are first translated to current level sequences using the $Q$-mer map, and then the (continuous) current level sequences are quantized to finite levels (*e.g.* three levels for $HQ3$) by hard thresholding the current levels.

##### 1.3 Accessing HQAlign on github

HQAlign requires python 3, and the installation guideline can be found on github.
The software is available at: `https://github.com/joshidhaivat/HQAlign.git`

```
usage:  python hqalign.py [-h] -r REF -i READS -o OUTPUT [-t THREADS] [-k
KMER]
```

arguments:

| | |
|---|---|
| `-h, --help` | show this help message and exit |
| `-r REF, --ref REF` | reference genome filename in fasta format |
| `-i READS, --reads READS` | directory location of read files in fasta format (with file extension `.fasta`) |
| `-o OUTPUT, --output OUTPUT` | location of directory of output files |
| `-t THREADS, --threads THREADS` | maximum number of parallel threads (default=4) |
| `-k KMER, --kmer KMER` | minimizer length for hybrid step (default=18) |

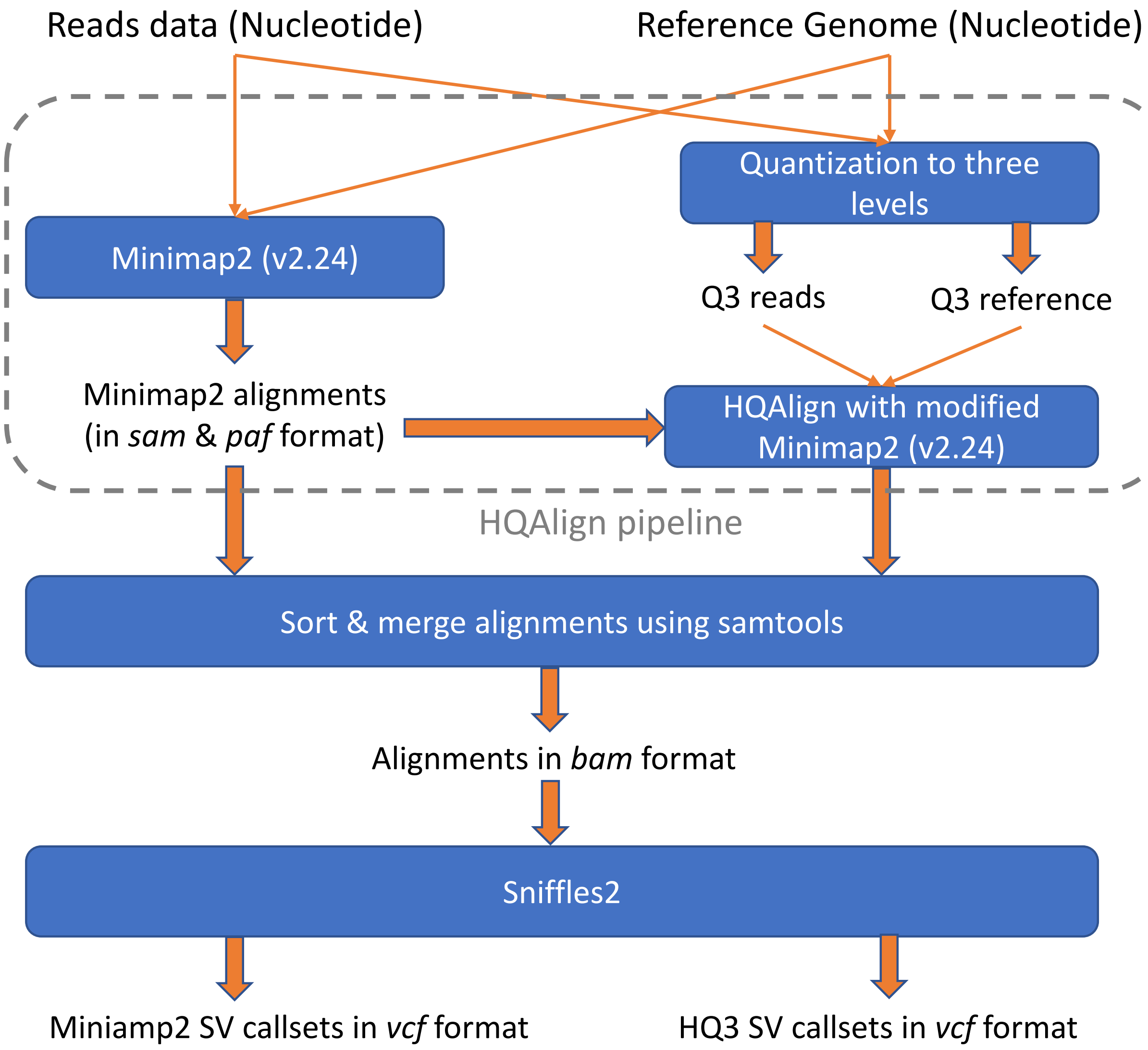


Figure 2: Complete pipeline for SV calling using minimap2 and HQAlign.

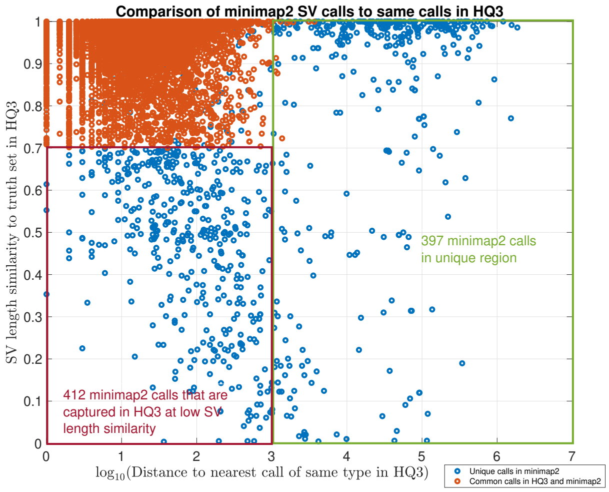


(a)

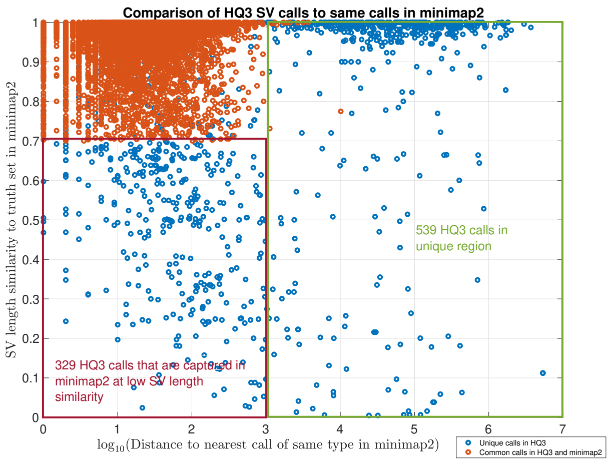


(b)

Figure 3: **Comparison of SV calls made by minimap2 and** $HQ3$ **to other method.** (a) For the complementary calls (in blue) and common calls (in red) made by minimap2, this figure compares SV length similarity and distance to nearest SV in HQ3 of the same type. 397 complementary calls made by minimap2 are in unique region, whereas 412 complementary calls in minimap2 are captured in neighboring region (within 1000 bp) in $HQ3$ but with a low SV length similarity. (b) For the complementary calls (in blue) and common calls (in red) made by $HQ3$, this figure compares SV length similarity and distance to nearest SV in minimap2 of the same type. 539 complementary calls made by $HQ3$ are in unique region, whereas 329 complementary calls in $HQ3$ are captured in neighboring region (within 1000 bp) in minimap2 but with a low SV length similarity.

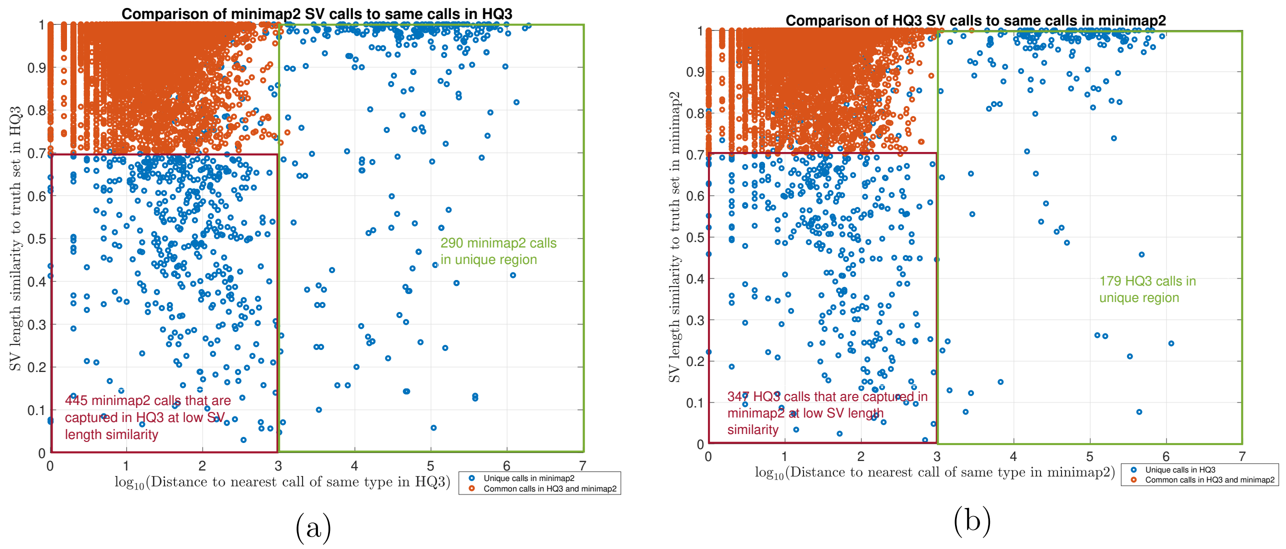


Figure 4: **Comparison of SV calls to HG002 truth set.** (a) For the complementary calls (in blue) and common calls (in red) made by minimap2, this figure compares SV length similarity and distance to nearest SV in HQ3 of the same type. 290 complementary calls made by minimap2 are in unique region, whereas 445 complementary calls in minimap2 are captured in neighboring region (within 1000 bp) in $HQ3$ but with a low SV length similarity. (b) For the complementary calls (in blue) and common calls (in red) made by $HQ3$, this figure compares SV length similarity and distance to nearest SV in minimap2 of the same type. 179 complementary calls made by $HQ3$ are in unique region, whereas 347 complementary calls in $HQ3$ are captured in neighboring region (within 1000 bp) in minimap2 but with a low SV length similarity.
